# Neuropilin 1 sequestration by neuropathogenic mutant glycyl-tRNA synthetase is permissive to vascular development and homeostasis

**DOI:** 10.1101/081513

**Authors:** James N. Sleigh, Adriana Gómez-Martín, Giampietro Schiavo

**Affiliations:** Sobell Department of Motor Neuroscience and Movement Disorders, Institute of Neurology, University College London, London WC1N 3BG, UK

**Keywords:** aminoacyl-tRNA synthetase (ARS), Charcot-Marie-Tooth disease type 2D (CMT2D), distal spinal muscular atrophy type V (dSMA-V), *GARS*, glycyl-tRNA synthetase (GlyRS), peripheral neuropathy

## Abstract

It remains a mystery how dominantly inherited mutations in the housekeeping gene *GARS*, which encodes glycyl-tRNA synthetase (GlyRS), mediate selective peripheral nerve toxicity resulting in the currently incurable Charcot-Marie-Tooth disease type 2D (CMT2D). A recent study identified the transmembrane receptor protein neuropilin 1 (Nrp1) as a substrate for aberrant extracellular binding of mutant GlyRS. Formation of the Nrp1/mutant GlyRS complex antagonises the interaction of Nrp1 with one of its main natural ligands, vascular endothelial growth factor-A (VEGF-A), contributing to neurodegeneration. Reduced binding of Nrp1 to VEGF-A is known to disrupt blood vessel development and growth. We therefore analysed capillary architecture at early and later symptomatic time points in CMT2D mouse muscles, retina, and sciatic nerve, as well as in embryonic hindbrain. Assessing capillary diameter, density, and branching, we observed no differences between wild-type and mutant animals from embryonic development to three months, spanning the duration over which numerous sensory and neuromuscular phenotypes manifest. This work indicates that mutant GlyRS-mediated disruption of Nrp1/VEGF-A signalling is permissive to capillary maturation and maintenance in CMT2D mice.

**Summary Statement:** Although the multi-functional neuropilin 1/VEGF-A signalling pathway is impaired by dominant pathogenic GlyRS variants, the vascular system remains unperturbed in *Gars* mutant mice.

## Introduction

*GARS* is a widely and constitutively expressed gene that encodes glycyl-tRNA synthase (GlyRS), which serves to covalently attach glycine to its transfer RNA (tRNA), priming it for protein translation (Antonellis and Green, 2008). Dominant mutations in *GARS* cause Charcot-Marie-Tooth disease type 2D (CMT2D, OMIM 601472), a disorder of the peripheral nerves with the principal clinical symptom of muscle wasting originating in the extremities (Antonellis et al., 2003). A battery of evidence indicates that these *GARS* mutations cause a toxic, gain-of-function (Grice et al., 2015; Motley et al., 2011; Nangle et al., 2007; Seburn et al., 2006; Xie et al., 2007), and that this is at least partially reliant upon the exposure to the protein surface of regions usually buried in wild-type GlyRS (He et al., 2011). Recently published work suggests that these neomorphic regions cause mutant GlyRS to aberrantly bind to the transmembrane receptor protein neuropilin 1 (NRP1) (He et al., 2015). NRP1 functions in both the nervous and vascular systems by binding to the extracellular signalling molecules semaphorin 3A (SEMA3A) and vascular endothelial growth factor-A 165 (VEGF-A_165_) via the co-receptors plexin A and VEGFR2, respectively (**Fig. S1A**) (Neufeld et al., 2002). In the study by Yang and colleagues, mutant GlyRS was shown to specifically interact with the extracellular region of NRP1 integral to VEGF-A_165_ binding (the b1 domain) (Gu et al., 2002), and thereby antagonise NRP1/VEGF-A signalling (**Fig. S1B**) (He et al., 2015). CMT2D embryonic day 13.5 (E13.5) hindbrains phenocopied the facial motor neuron migration defects of mice deficient in VEGF-A_164_ (murine equivalent of human VEGF-A_165_) and Nrp1 (Schwarz et al., 2004). Moreover, genetically reducing Nrp1 levels enhanced the severity of CMT2D pathology, whereas providing additional VEGF-A_165_ via lentiviral intramuscular injections improved the phenotype.

VEGF-A functions in a diverse array of neuronal processes (Mackenzie and Ruhrberg, 2012; Ruiz de Almodovar et al., 2009), and a mild reduction in VEGF-A causes progressive motor neuron degeneration in mice (Oosthuyse et al., 2001), partially explaining why antagonisation of Nrp1/VEGF-A signalling could be contributing to the neuronal pathology in CMT2D. However, NRP1 and VEGF-A are also critical to blood vessel formation (vasculogenesis), development/expansion (angiogenesis), and growth (arteriogenesis) (Kofler and Simons, 2015; Plein et al., 2014). Homozygous *Nrp-1* (Gerhardt et al., 2003; Kawasaki et al., 1999), homozygous *VEGFR2* (Shalaby et al., 1995), and heterozygous *VEGF-A* (Carmeliet et al., 1996; Ferrara et al., 1996) null mice all display severe embryonic vascular deficits that greatly contribute to embryonic lethality. Though mutating residues in the VEGF-A binding site of *Nrp1* does not phenocopy the severity of the knockout mice (Fantin et al., 2014; Gelfand et al., 2014), it reduces blood vessel branching in the embryonic hindbrain and causes a post-natal impairment in angiogenesis and arteriogenesis (Fantin et al., 2014), resembling the milder phenotype of mice lacking Nrp1-binding VEGF-A isoforms (Ruhrberg et al., 2002). This body of work suggests that Nrp1 has both VEGF-dependent and VEGF-independent functions in blood vessels, and that Nrp1/VEGF-A signalling is perhaps more important for vascularisation post- than pre-birth (Fantin et al., 2014). Consistent with this, blocking VEGF-A binding to Nrp1 with a monoclonal antibody impairs vascular remodeling in the peri-natal mouse retina (Pan et al., 2007).

Although the exact function of VEGF-A signalling through Nrp1 in blood vessels has not been fully elucidated at all stages of life, its central importance to the vascular system is absolutely clear. Given that mutant GlyRS competes with VEGF-A for Nrp1 binding, and that GlyRS is secreted and found in circulating serum of humans and mice (Grice et al., 2015; He et al., 2015; Park et al., 2012), we decided to assess the effect of mutant GlyRS on the architecture of capillaries in a murine model of CMT2D.

## Results and Discussion

Nrp1 is widely expressed in a range of different tissues (Soker et al., 1998), localising to the vascular system (Fantin et al., 2011; Fantin et al., 2010) and many neuronal types, including motor neurons (Feldner et al., 2005; Huber et al., 2005; Moret et al., 2007). As mutated GlyRS binds to Nrp1 impacting the motor nervous system of *Gars* mice (He et al., 2015), and CMT2D is a neuromuscular condition, we wanted to assess the expression pattern of Nrp1 in a post-natal tissue with both vasculature and motor nerves. We therefore performed immunohistochemistry on wholemount distal, fast-twitch lumbrical and proximal, slow-twitch transversus abdominis (TVA) muscles from one and three month old wild-type mice. These muscles are thin and flat, permitting the visualisation of the entire vascular and nervous systems (Murray et al., 2014; Sleigh et al., 2014a). In these muscles at both time points, Nrp1 (white) did not specifically localise to SV2/2H3^+^ (green) motor neurons or S100^+^ (blue) Schwann cells (**Fig. 1A-B**). Rather, its expression consistently coincided with the endothelium-binding isolectin B_4_ (IB4, red) (**Fig. 1C, S2A-B**). Nrp1 could be observed faintly surrounding the motor neurons contiguous with IB4, but peripheral to the staining of both S100 and myelin basic protein (Mbp, yellow) (**Fig. 1C, S2C**). These results indicate that Nrp1 predominantly localises to the vascular system in post-natal skeletal muscle; however, we cannot exclude there being a low level of motor neuronal expression. Indeed, a previous report suggests that Nrp1 is found in motor axons of sectioned, adult gastrocnemius muscle (Venkova et al., 2014). The fundamental requirement for Nrp1 in motor nervous system development is undisputed (Feldner et al., 2005; Huber et al., 2005; Huettl et al., 2011; Moret et al., 2007), but its post-natal function and localisation remains less well defined. Motor neuron-specific ablation of Nrp1 using the *Olig2* promoter is reported to cause post-natal motor axon loss and muscle atrophy (Helmbrecht et al., 2015), but Nrp1 was deleted developmentally (*Olig2* is expressed from ≈E8.5 (Lu et al., 2000)) rather than at or post-birth. It is possible that low levels of post-natal motor neuronal Nrp1 expression provide sufficient substrate for mutant GlyRS to contribute to peripheral nerve degeneration in CMT2D mice. However, given that developmental defects are observed in *Gars* mice (He et al., 2015; Sleigh et al., 2016 preprint), and that Nrp1 expression is developmentally downregulated in neuronal tissue (Bovenkamp et al., 2004; Fujisawa et al., 1995), we should not rule out that mutant GlyRS-mediated inhibition of Nrp1/VEGF-A signalling during development may be predisposing the motor system to the subsequent degeneration observed at later, post-natal stages (Sleigh et al., 2014b; Spaulding et al., 2016).

**Figure 1.**
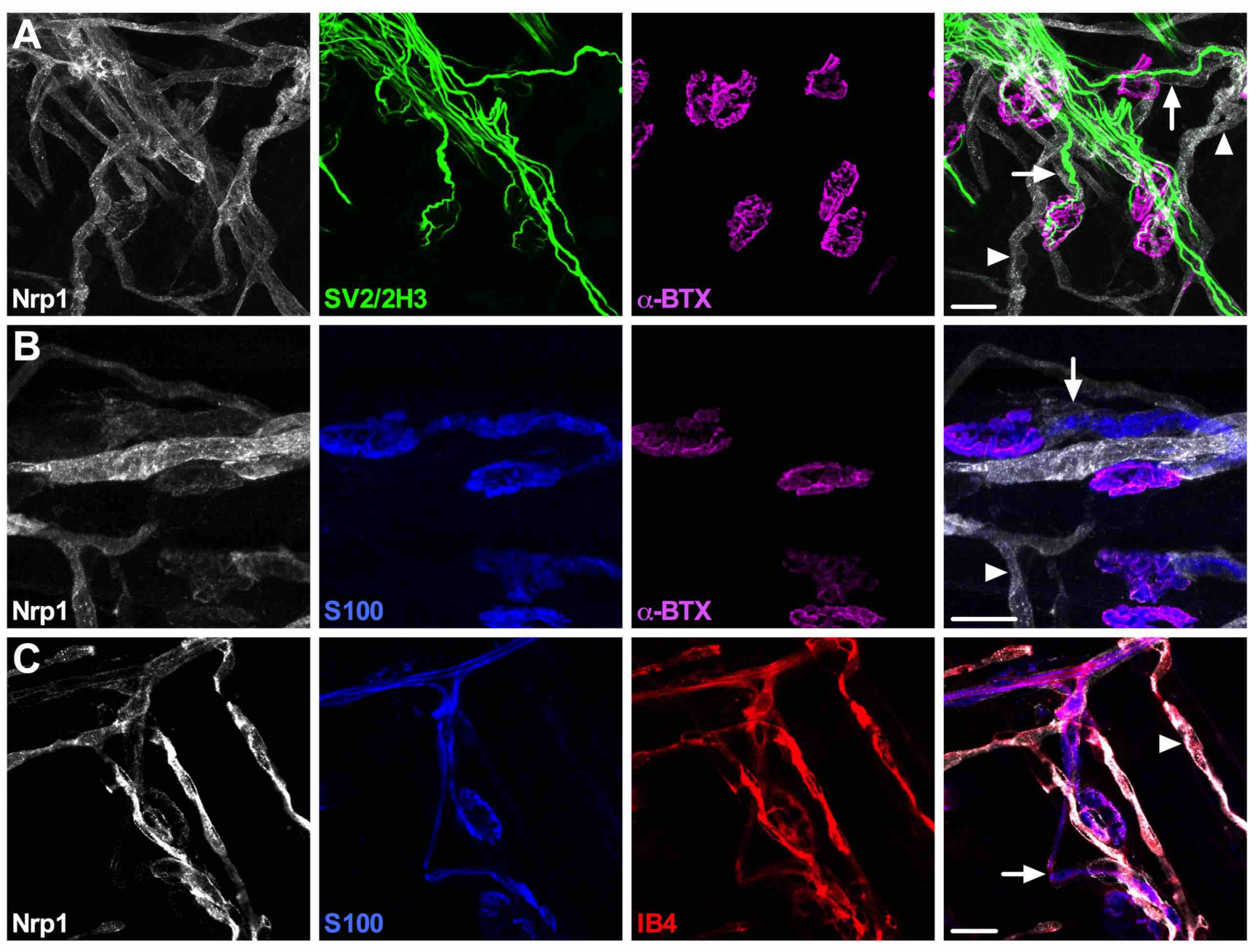
Nrp1 localises to skeletal muscle blood vessels but not motor neurons. Representative Nrp1 staining in one month-old wild-type lumbrical muscles. (**A**) Nrp1 (white) localises to structures surrounding, but not contiguous with, lower motor neurons (green, SV2/2H3), and in SV2/2H3^-^ areas. α-bungarotoxin (α-BTX, magenta) identifies post-synaptic acetylcholine receptors at the neuromuscular junction. (**B**) Nrp1 does not perfectly overlap with the Schwann cell marker S100 (blue) either. This was confirmed with a second glial cell marker, myelin basic protein (Mbp, Fig. S2C). (**C**) Nrp1 co-localises with IB4 (red), a lectin that labels endothelial cells, indicating that Nrp1 is found in muscle capillaries. For all images, arrows highlight Nrp1 staining associated with motor neurons/myelin, while arrowheads indicate Nrp1^+^ blood vessels. A similar pattern of staining was observed at three months and in the TVA muscle at both time points (data not shown). A-B are collapsed Z-stack images and C is a single plane image. Scale bars = 20 μm.

Irrespective of the importance of Nrp1 in motor neurons, it is clear that it is highly expressed in blood vessels embedded in skeletal muscle. We therefore decided to assess structural features of the muscle vascular bed in wild-type and *Gars*^*C201R/+*^ mice using IB4 staining (**Fig. S3**). *Gars*^*C201R/+*^ mice display a range of sensory and motor defects modelling CMT2D (Achilli et al., 2009; Sleigh et al., 2016 preprint; Sleigh et al., 2014b; Spaulding et al., 2016). Confocal Z-stack images were taken throughout the entire depth of one and three month old lumbrical and TVA muscles (**Fig. 2A,B**), and capillary diameter (**Fig. 2C,D**), density (**Fig. 2E,F**), and branching (**Fig. 2G,H**) assessed. These time points represent early and later symptomatic stages of disease in *Gars*^*C201R/+*^ mice, and were chosen to complement previously performed in-depth motor and sensory phenotypic analyses (Sleigh et al., 2016 preprint; Sleigh et al., 2014b). We saw no significant difference between wild-type and mutant *Gars* capillary beds in any of the parameters analysed, suggesting that *Gars*^*C201R/+*^ vasculature is unimpaired in fast and slow twitch skeletal muscles. The capillary diameter was consistent across muscles and varied little over time, while capillary and branching densities showed variability between muscles, and appeared to decline with age, presumably due to muscle growth. These differences confirm the suitability of this strategy for detecting changes in the vasculature induced over time. We also stained muscles for Nrp1 and VEGFR2, but found no obvious discrepancies in the level of expression or localisation between genotypes (**Fig. S4**).

**Figure 2.**
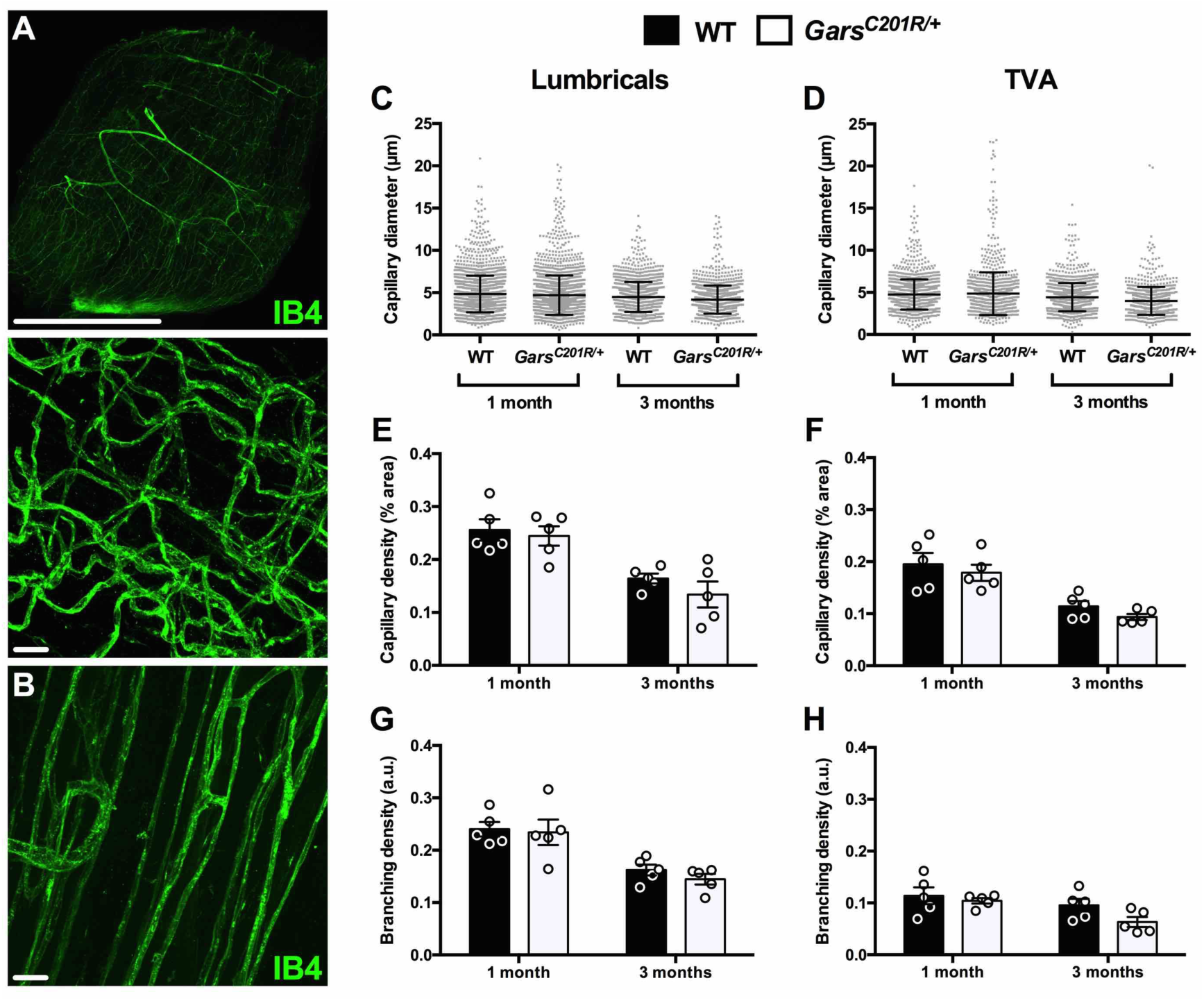
Capillaries appear unaffected in *Gars*^*C201R/+*^ skeletal muscles. (**A-B**) Representative single plane (A, top) and collapsed Z-stack (A, bottom and B) images of IB4 staining in one month lumbrical (A) and TVA muscles (B). Scale bars = 1 mm (A, top) and 20 μm (A, bottom and B). (**C, E, G**) No defects in capillary diameter (C, age, *P* = 0.007; genotype, *P* = 0.090; interaction, *P* = 0.508), density (E, age, *P* < 0.001; genotype, *P* = 0.290; interaction, *P* = 0.625), or branching density (F, age, *P* < 0.001; genotype, *P* = 0.468; interaction, *P* = 0.721) were observed in *Gars*^*C201R/+*^ mouse lumbrical muscles at one and three months. (**D, F, H**) The TVA muscle also showed no difference in capillary diameter (D, age, *P* = 0.875; genotype, *P* = 0.264; interaction, *P* = 0.359), density (F, age, *P* < 0.001; genotype, *P* = 0.223; interaction, *P* = 0.889), or branching density (H, age, *P* = 0.020; genotype, *P* = 0.086; interaction, *P*= 0.340). All data sets were analysed with two-way ANOVAs. *n* = 5.

Nrp1 is indispensable for vascularisation of the mouse retina and central nervous system, but is thought to be less critical for the vasculature of other tissues such as muscle (Tata et al., 2015). *In vivo* disruption of Nrp1 binding to VEGF-1 has previously been shown to disturb post-natal vascular density and patterning of the retina at 21 days, and vessel branching in embryonic hindbrains (Fantin et al., 2014; Gelfand et al., 2014). We therefore assessed the IB4^+^ capillary network in one and three month old retinas (**Fig. 3**), and E13.5 hindbrains (**Fig. 4A,C-E**). *Gars* mice had similar retinal capillary diameters (**Fig. 3B**), densities (**Fig. 3C**), and branching (**Fig. 3D**) as wild-type at both time points. There was also no difference in the number of major radial branches emanating from the central retina (**Fig. 3E**). The mutant hindbrain equally showed no distinctions in blood vessel structures between genotypes (**Fig. 4C-E**). Similar to skeletal muscles, no obvious differences in Nrp1 and VEGFR2 staining in either retinas or hindbrains were seen (data not shown), which is consistent with western blotting showing that Nrp1 and VEGFR2 protein levels remain unaltered in embryonic *Gars* neural tissues (He et al., 2015). Finally, we assessed vascular density in sectioned sciatic nerves from one month old mice, in order to see whether post-natal neuronal tissue was affected. Anti-platelet endothelial cell adhesion molecule 1 (Pecam1) was used instead of IB4 due to superior staining of the vasculature in this tissue (**Fig. 4B**). Mutant mice showed a non-significant trend towards increased capillary density (**Fig. 4F**); however, this is likely simply caused by the reduced axon calibres of mutant mice (Achilli et al., 2009).

**Figure 3.**
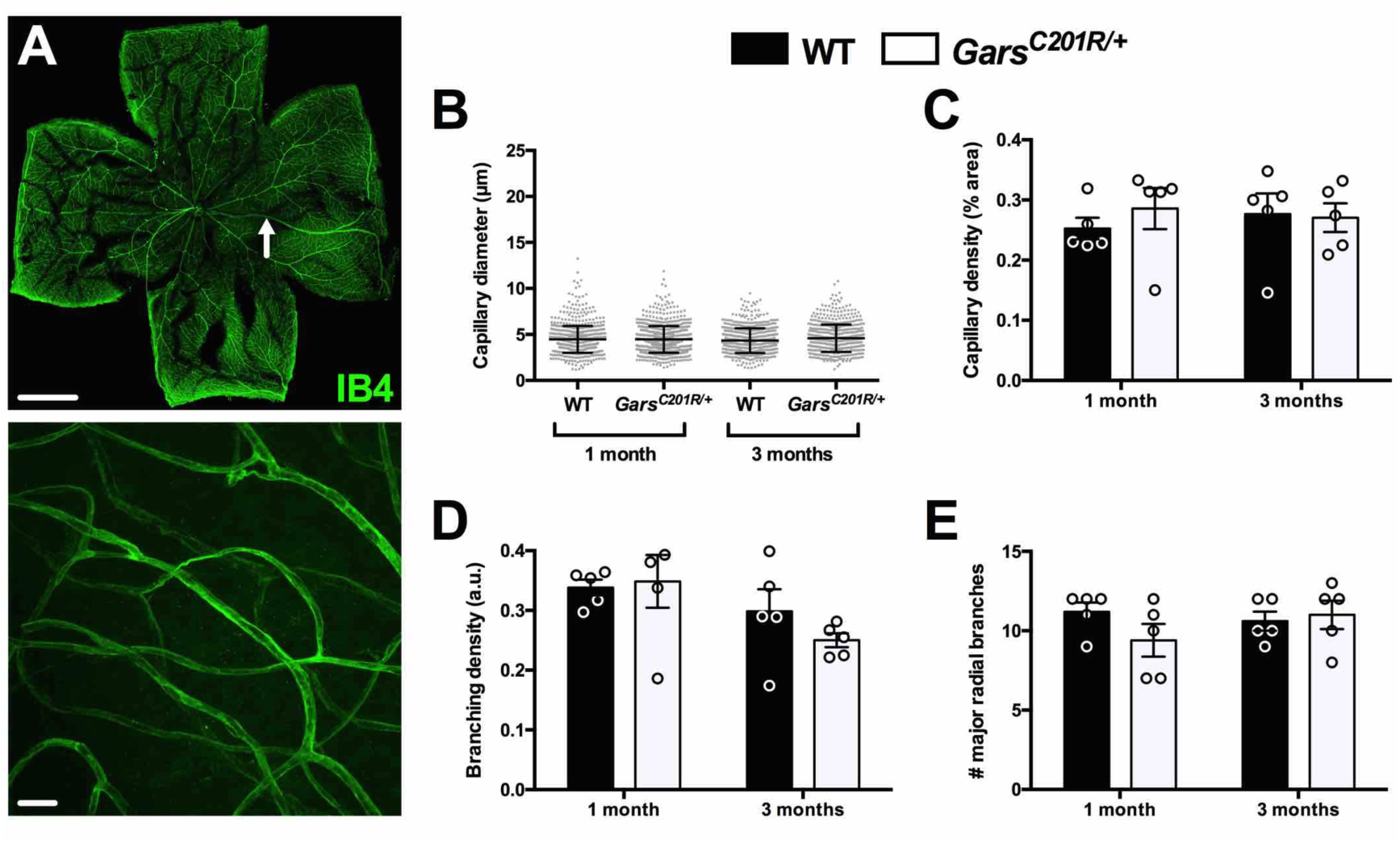
The capillary network is unaffected in *Gars*^*C201R/+*^ retinas. (**A**) Representative single plane (top) and collapsed Z-stack (bottom) images of IB4 staining in one month retina. Scale bars = 1 mm (top) and 20 μm (bottom). (**B-E**) No defects in capillary diameter (B, age, *P* = 0.823; genotype, *P* = 0.112; interaction, *P* = 0.105), density (C, age, *P* = 0.878; genotype, *P* = 0.638; interaction, *P* = 0.504), branching density (D, age, *P* = 0.042; genotype, *P* = 0.548; interaction, *P* = 0.345), or major radial branch (arrow in A) number (E, age, *P* = 0.541; genotype, *P* = 0.395; interaction, *P* = 0.188) were observed in *Gars*^*C201R/+*^ mouse retinas at one and three months. All data sets were analysed with two-way ANOVAs. *n* = 5.

**Figure 4.**
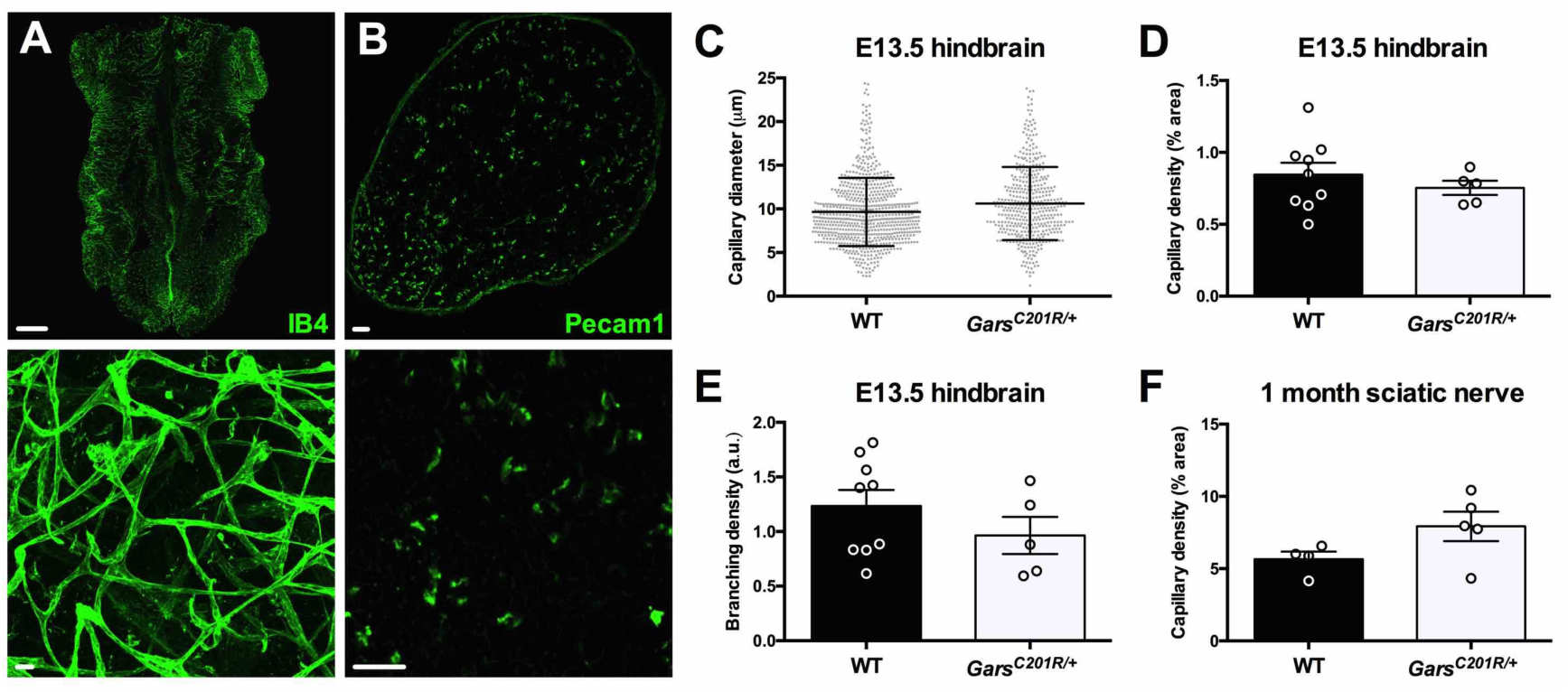
*Gars* mutant embryonic hindbrains and sciatic nerves show no vascular defects. (**A-B**) Representative single plane (A, top and B) and collapsed Z-stack (A, bottom) images of IB4 and Pecam1 staining in E13.5 hindbrain (A) and one month sciatic nerve (B), respectively. Scale bars = 1 mm (A, top) and 20 μm (A, bottom, and B). (**C-E**) Capillary diameter (C, *P* = 0.593), density (D, *P* = 0.456), and branching density (E, *P* = 0.278) are all unimpaired in *Gars*^*C201R/+*^ E13.5 hindbrains. *n* = 9 wild-type and 5 *Gars*^*C201R/+*^. (**F**) Mutant sciatic nerves show no difference in capillary density at one month (*P* = 0.111). *n* = 4 wild-type and 5 *Gars*^*C201R/+*^. All data were analysed with unpaired, two-tailed *t*-tests.

Mutant GlyRS appears to compete with VEGF-A for extracellular binding to the transmembrane receptor protein Nrp1 contributing to the peripheral nerve pathology observed in CMT2D (He et al., 2015). Reduced binding of VEGF-A to Nrp1 has previously been shown to impair post-natal angiogenesis and arteriogenesis (Fantin et al., 2014; Gelfand et al., 2014). We thus set out to determine whether the capillary network of mutant *Gars* mice is altered either in the post-natal period or developmentally. We found that Nrp1 was highly expressed in post-natal skeletal muscle blood vessels, but not in motor neurons (**Fig. 1**); however, capillary diameter, density, and branching were unaffected by mutant GlyRS in both distal and proximal muscles (**Fig. 2**). This was replicated in adult retinas (**Fig. 3**), one month sciatic nerves (**Fig. 4F**), and embryonic hindbrains (**Fig. 4C-E**). We have therefore shown that the vascular system is unaffected in *Gars*^*C201R/+*^ mice from embryonic development to adulthood. Extra-neuronal tissue pathology and disease mechanisms have been reported in mouse models of a number of peripheral nerve conditions including spinal muscular atrophy (Sleigh et al., 2011), Kennedy’s disease (Cortes et al., 2014; Lieberman et al., 2014), and amyotrophic lateral sclerosis (Puentes et al., 2016), appearing to mirror the human conditions, at least in some of their most debilitating incarnations (Somers et al., 2016). Nevertheless, we have convincingly ruled out that non-neuronal pathology extends to CMT2D, which is in keeping with the clinical presentation of patients.

So why does antagonisation of Nrp1/VEGF-A signalling affect the peripheral nervous system, but not the vascular system, when it is critical for the functioning of both? First of all, this could reflect discrepancies between the mild *Gars*^*C201R/+*^ model used here and the more severe *Gars*^*Nmf249/+*^ mouse used by He *et al.* (2015). It is possible that antagonisation of Nrp1/VEGF-A signalling *in vivo* is specific to the *Gars*^*Nmf249/+*^ mutant; however, this is unlikely because all CMT2D-associated *GARS* mutations tested to date aberrantly bind to Nrp1 *in vitro*, although C201R has not been verified. Instead, timing and location of Nrp1 and GlyRS expression could conspire to ensure neuronal specificity. GlyRS is secreted and present in serum, but it may not be found in the immediate vicinity of Nrp1 on blood vessels. For instance, if GlyRS is secreted by terminal Schwann cells at the neuromuscular synapse, GlyRS may only gain access to Nrp1 on motor nerve terminals. Alternatively, mutant GlyRS could compete with VEGF-A for Nrp1 binding while simultaneously impinging upon additional, VEGF-independent pathways. Finally, the lack of a *Gars* vascular phenotype may simply reflect that a mild dampening of Nrp1/VEGF-A signalling preferentially impacts the neuronal-specific functions of the pathway. This scenario is perhaps the most likely, as it corroborates the previous observation that VEGF-A-deficient mice display selective degeneration of motor neurons (Oosthuyse et al., 2001).

In summary, we have clarified that CMT2D mice display a strict neuropathology, and that GlyRS-mediated disruption of Nrp1/Vegf-A signalling appears to be permissive to capillary maturation and maintenance. This indicates that there is a possible concentration-dependent dual action of the Nrp1/VEGF pathway in both nervous and vascular systems.

## Materials and Methods

### Animals

*Gars*^*C201R/+*^ mice were maintained as heterozygote breeding pairs on a predominantly C57BL/6 background and genotyped as described previously (Achilli et al., 2009). Mice sacrificed at one and three month time points were 28-34 and 89-97 days old, respectively. Multiple tissues were simultaneously harvested from both males and females. Mouse handling and experiments were performed under license from the UK Home Office in accordance with the Animals (Scientific Procedures) Act (1986), and approved by the University College London – Institute of Neurology Ethics Committee.

### Tissue preparation and immunohistochemistry

All steps were performed at room temperature, apart from overnight incubations conducted at 4°C. Lumbrical and TVA muscles dissected and immunohistochemically labelled as previously described (Murray et al., 2014; Sleigh et al., 2014a; Sleigh et al., 2014b). Eyes were removed and fixed in 4% (w/v) paraformaldehyde (PFA, Electron Microscopy Sciences, Hatfield, PA) for 2 h, before retinas were dissected and stained as reported previously (Pitulescu et al., 2010). Sciatic nerves were dissected from mice transcardially perfused with 4% PFA, post-fixed for 2 h, and processed and 10 μm section stained as previously described (Sleigh et al., 2016 preprint). E13.5 hindbrains were dissected and stained using published protocols (Fantin et al., 2013). All tissues were incubated with primary antibodies overnight (**Table S1**), and sometimes with 1 mg/ml isolectin B_4_ (IB4) biotin conjugate (Sigma Aldrich, St. Louis, MO, L2140) in phosphate buffered saline (PBS, 137 mM NaCl [Sigma, S3014], 10 mM Na_2_HPO_4_ [Sigma, S3264], 2.7 mM KCl [Sigma, P9541], 1.8 mM KH_2_PO_4_ [Sigma, P9791]) at 1/250. The following day, combinations of Alexa Fluor-labelled secondary antibodies (Life Technologies, Carlsbad, CA, A-21202, A-21432, A-21447, A-31570, A-31572) at 1/250, 2 mg/ml streptavidin Alexa Fluor 488 conjugate (Life Technologies, S-11223) at 1/250, and fluorophore-conjugated α-bungarotoxin (α-BTX, Life Technologies, B-13422 and B-35451; Biotium, Fremont, CA, 00002) at 1/1000 in PBS were used for 2 h, before the tissues were washed and mounted in fluorescent mounting medium (Dako, Glostrup Municipality, Denmark, S3023) for imaging.

### Imaging and analysis

Tissues were imaged using a LSM 780 laser scanning microscope (Zeiss, Oberkochen, Germany) and images analysed using ImageJ software (https://imagej.nih.gov/ij/). All samples were imaged and analysed blinded to genotype. No samples were excluded from the analyses once imaged. IB4-stained muscles, retinas, and hindbrains, and Pecam1-stained sciatic nerve sections were used for capillary analyses as performed previously (Somers et al., 2016; Somers et al., 2012). Maximal Z-resolution stacked images across the entire depth of wholemount tissues were obtained at 40x (for muscles) and 20x (for retinas and hindbrains) magnification and 1024x1024 pixel resolution. Sciatic nerve sections were single plane images taken at 20x and 2812x2812 pixel resolution. Three non-overlapping fields of view were imaged per lumbrical muscle, and four lumbricals analysed per animal (12 Z-stacks in total). Eight non-redundant Z-stack images were obtained per TVA and retina, and four per hindbrain, and analysed for each animal. Similar tissue regions were imaged across samples. Five different sections were analysed per sciatic nerve. For measuring capillary diameters, a uniform grid (area per point: 500 μm^2^) was overlaid onto 3D-projected (Max Intensity) images, and every capillary found at a grid intersection measured (except those that were branching), with repeated measures of the same capillary accepted (**Fig. S3A-B**). Over 400 capillary diameters were measured and averaged per tissue per genotype. For capillary density analyses, images were 3D-projected (Max Intensity), converted to binary (capillaries assigned to black), and particles analysed (**Fig. S3A, C**). The summed black particle number was then divided by the number of sections to account for Z-stack depth. To assess branching density, capillary bifurcations were counted using the Cell Counter plugin on projected images and the total branches divided by the number of sections (**Fig. S3A, D**). Non-projected image stacks were used to ensure the counted branches were bifurcations and not crossing capillaries.

### Statistical analysis

Data were assumed to be normally distributed unless evidence to the contrary could be provided by the D'Agostino and Pearson omnibus normality test. GraphPad Prism 5 (La Jolla, CA) was used for all analyses. Means ± standard error of the mean are plotted, except for individual capillary diameter data, which represent mean and standard deviation. All analyses were performed on animal mean values.

## Author Contributions

J.N.S. conceived the work; J.N.S., A.G.-M., and G.S. designed the experiments; J.N.S. and A.G.-M. performed the experiments and analysed the data; all authors contributed to the writing of the paper and have approved submission of this work. The funders had no role in study design, data collection and analysis, decision to publish, or preparation of the manuscript.

## Acknowledgements

The authors would like to thank members of the Schiavo and Linda Greensmith (Institute of Neurology, UCL) laboratories for productive discussions, and Sergey S. Novoselov for experimental expertise.

## Abbreviations

α-BTX: α-bungarotoxin
CMTD2: Charcot-Marie-Tooth disease type 2D
DSHB: Developmental Studies Hybridoma Bank
GARS: GlyRS gene
GlyRS: glycyl-tRNA synthetase
IB4: isolectin B_4_
Mbp: myelin basic protein
Nrp1: neuropilin 1
PBS: phosphate buffered saline
Pecam1: platelet endothelial cell adhesion molecule 1
PFA: paraformaldehyde
SEMA3A: semaphorin 3A
SV2: synaptic vesicle 2
tRNA: transfer RNA
TVA: transversus abdominis
VEGF-A: vascular endothelial growth factor A
VEGFR2: vascular endothelial growth factor receptor 2
2H3: neurofilament

## Supplementary Figure Legends

**Figure S1.**
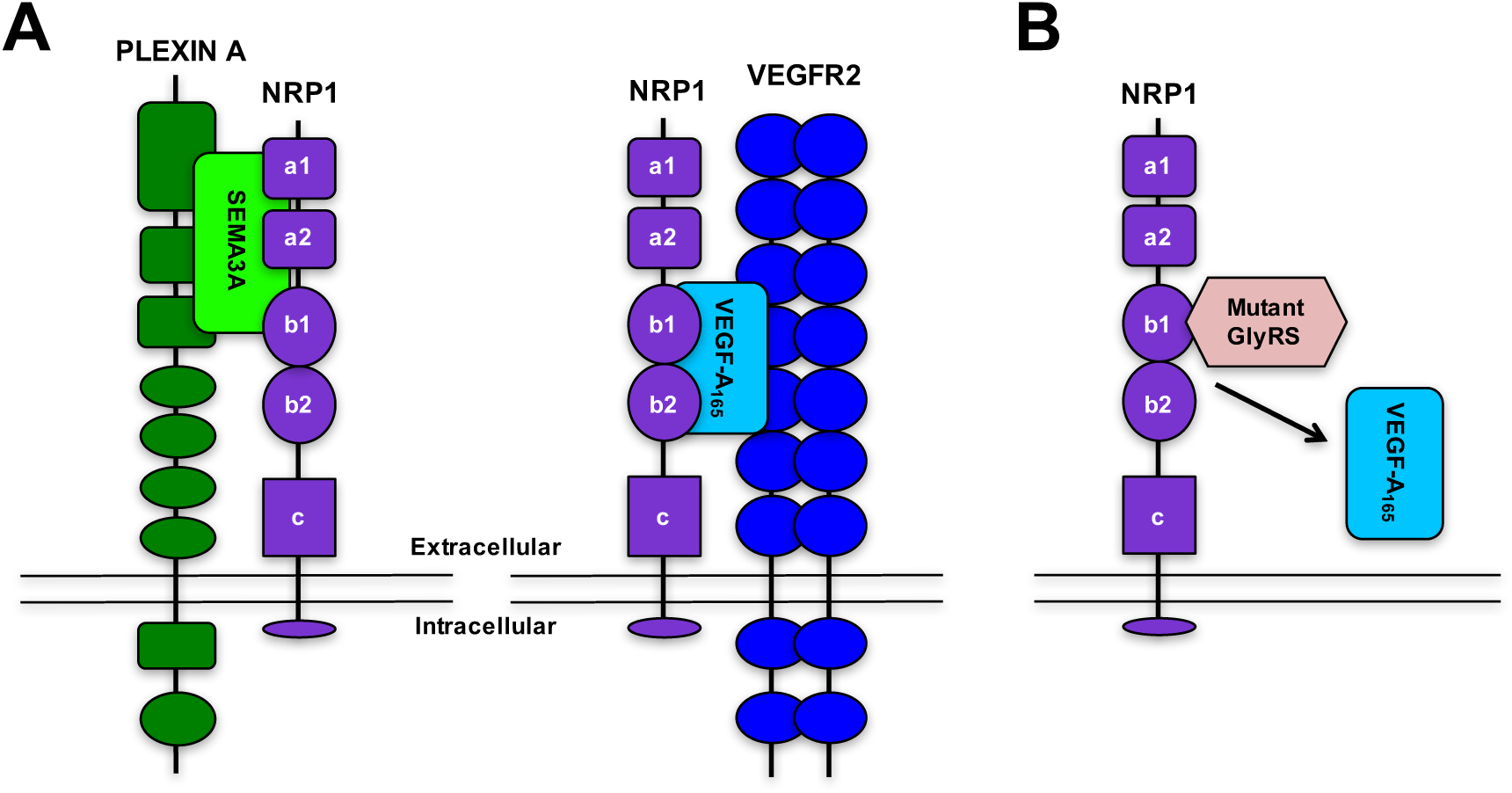
NRP1 is a membrane-bound receptor protein with diverse functions. (**A**) Schematic of NRP1 and two of its principal co-receptor proteins, PLEXIN A and VEGFR2, which bind to secreted glycoprotein ligands, SEMA3A and VEGF-A_165_, respectively (Neufeld et al., 2002). Binding of SEMA3A to the NRP1-PLEXIN A co-receptor complex is involved in several diverse biological processes (Epstein et al., 2015; Gu and Giraudo, 2013), the best characterised of which is guidance of growing axons (Fujisawa, 2004). VEGF-A_165_ and NRP1 function in vasculogenesis, angiogenesis, and arteriogenesis (Kofler and Simons, 2015), as well as in the nervous system (Mackenzie and Ruhrberg, 2012). SEMA3A and VEGF-A_165_ interact with different regions of NRP1; the a1 and a2 domains of NRP1 are essential for SEMA3A binding, while VEGF-A_165_ binding is principally dependent on the NRP1 b1 domain (Gu et al., 2002). (**B**) CMT2D-associated mutations in *GARS* cause a conformational opening of GlyRS, permitting the aberrant binding of mutant GlyRS to the b1 domain of NRP1. This competitively antagonises NRP1-VEGF-A signalling, which contributes to motor deficits observed in CMT2D mice. Schematics are not drawn to scale and are adapted from Plein *et al.* 2014 and He *et al.* 2015.

**Figure S2.**
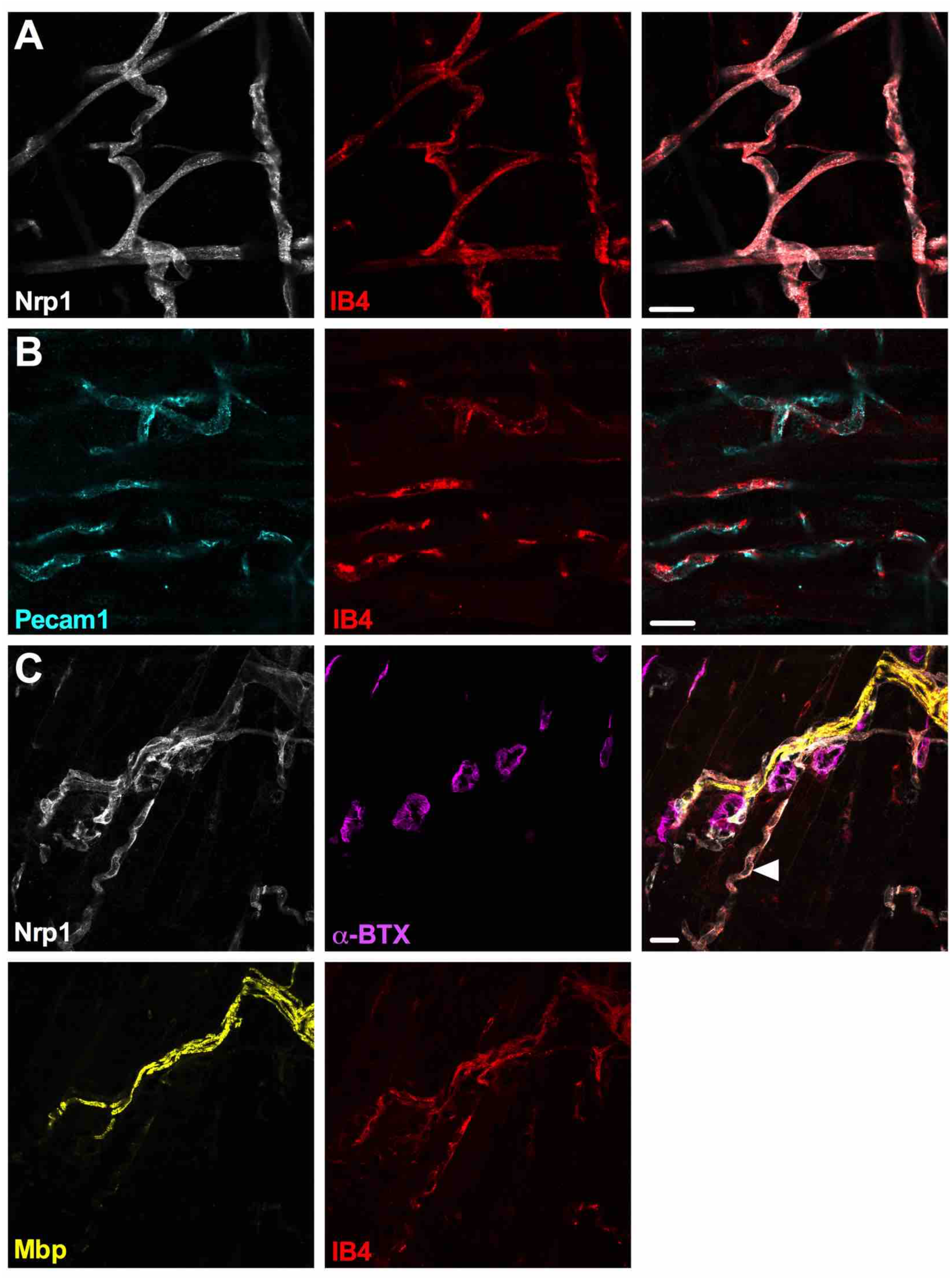
Nrp1 is expressed in IB4^+^ structures contiguous with the essential endothelium protein Pecam1. Representative images of Nrp1 localising to capillaries in one month-old wholemount lumbrical muscles. (**A**) Nrp1 (white) displays a high degree of co-localisation with IB4 (red), which marks the endothelium. (**B**) IB4 co-localises with platelet endothelial cell adhesion molecule 1 (Pecam1, cyan), an essential component of the endothelium. (**C**) Nrp1 localises to regions associated with the myelin marker protein Mbp (myelin basic protein, yellow, arrow), but is not perfectly contiguous with all glial cells. The arrowhead highlights a Nrp1^+^ capillary. Similar staining was observed in the TVA muscle (data not shown). All panels are single plane images. Scale bars = 20 μm.

**Figure S3.**
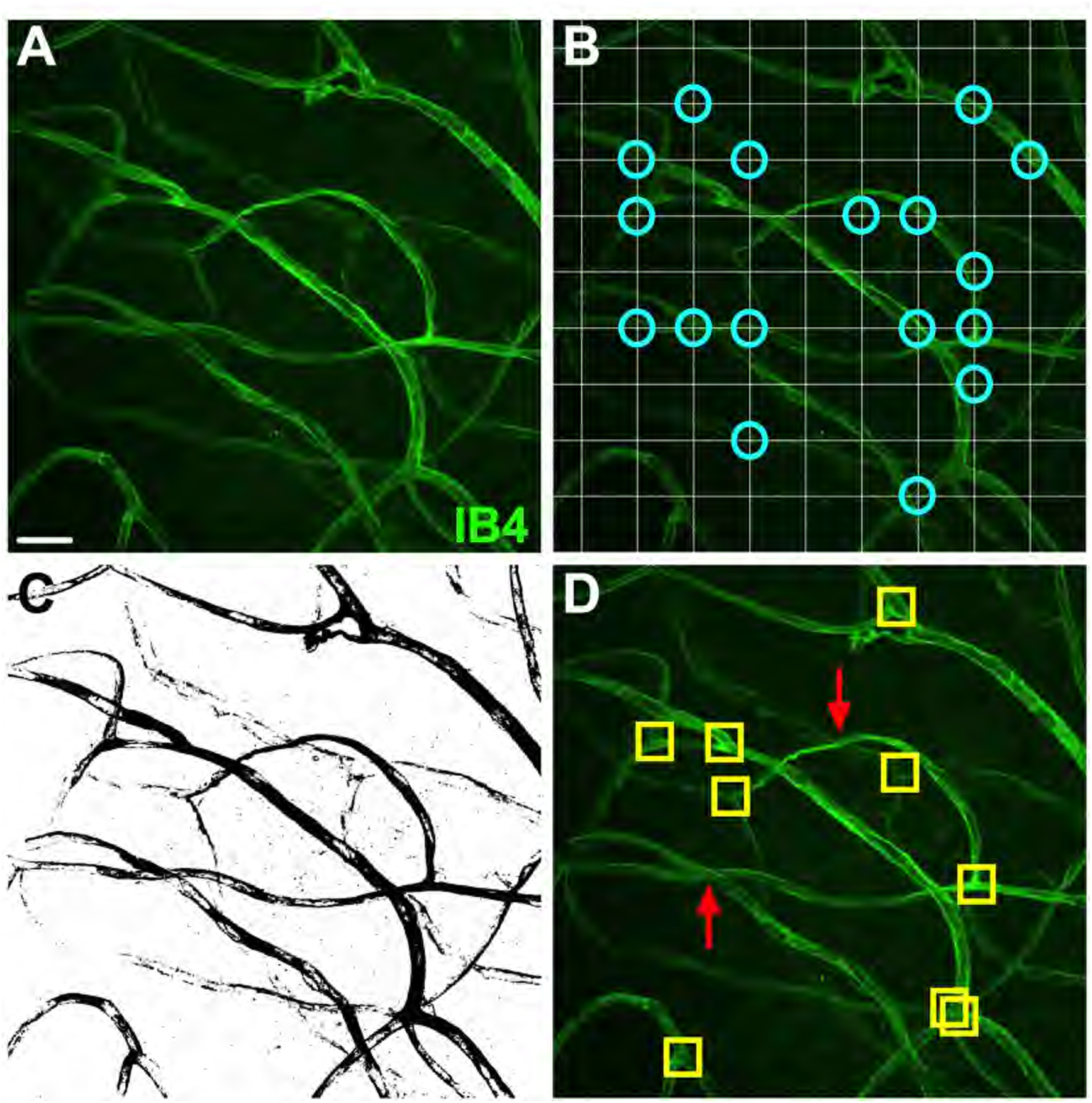
Capillary analyses. (**A**) Representative collapsed Z-stack image of IB4 staining in one month retina. Scale bar = 20 μm and applies to all images. (**B**) To measure capillary diameters, a uniform grid (white) was projected onto collapsed Z-stack images, and all capillary diameters found at grid intersections measured (cyan circles). (**C**) Collapsed Z-stacks were converted to binary and the sum of all black particles calculated using the Analyze Particles tool in order to measure capillary density. (**D**) Branching density was assessed using the Cell Counter plugin to mark all capillary bifurcations (yellow squares). Non-collapsed stacks were simultaneously used to ensure that crossing capillaries in different planes were not included in the count (red arrow examples).

**Figure S4.**
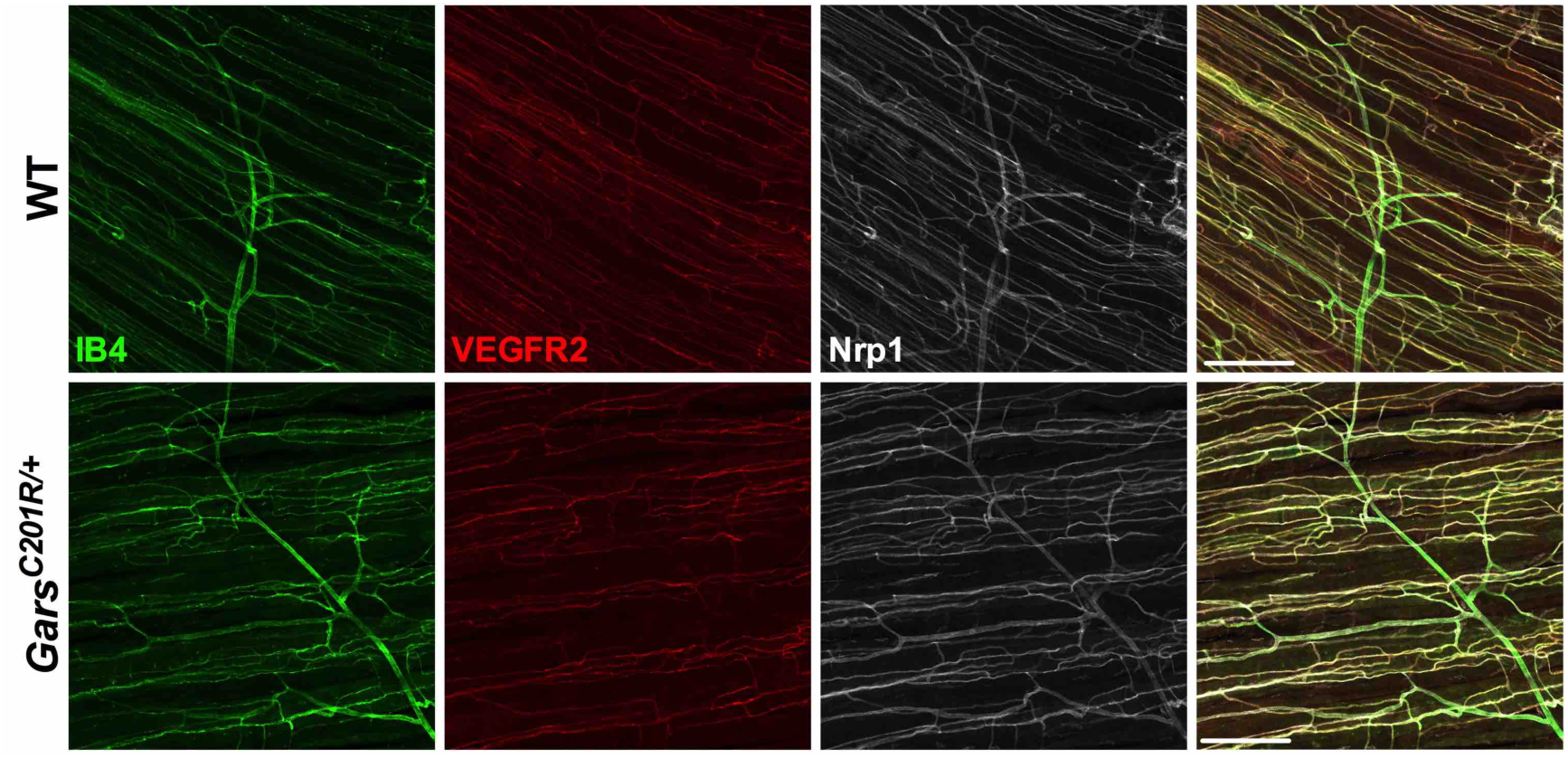
There are no ostensible distinctions in Vegfr2 and Nrp1 staining between wild-type and *Gars*^*C201R/+*^ skeletal muscles. (**A**) Representative Vegfr2 (red) and Nrp1 (white) staining in one month-old wholemount wild-type and *Gars*^*C201R/+*^ TVA muscles. Vegfr2 and Nrp1 colocalise with the endothelium marker IB4 (green) in both genotypes, and show no obvious differences in expression. A similar pattern was seen at one month and in the lumbrical muscles at both time points (data not shown). Both panels are single plane images. Scale bars = 200 μm.

## Supplementary Tables

**Supplementary Table 1.**
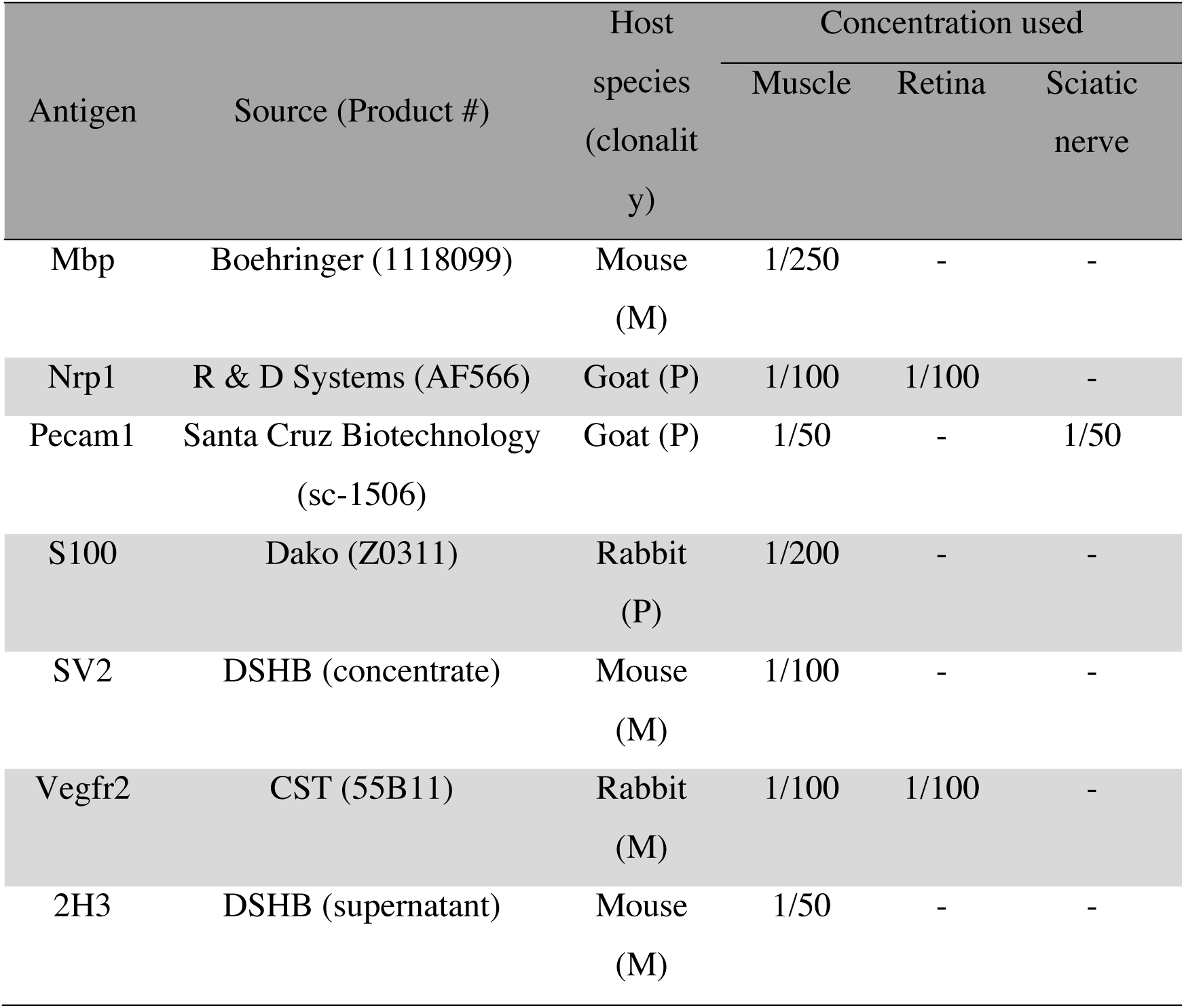
Primary antibodies used in the study. Mbp, myelin basic protein; Nrp1, neuropilin 1; Pecam1, platelet endothelial cell adhesion molecule 1; SV2, synaptic vesicle 2 (developed by Kathleen M. Buckley); Vegfr2, vascular endothelial growth factor receptor 2; 2H3, neurofilament (developed by Thomas M. Jessell and Jane Dodd). Boehringer, Ingelheim am Rhein, Germany; R & D Systems, Minneapolis, MN; Santa Cruz Biotechnology, Dallas, TX; Developmental Studies Hybridoma Bank (DSHB), Iowa City, IA; Cell Signalling Technology (CST), Danvers, MA. M, monoclonal; P, polyclonal.

| Antigen | Source (Product #) | Host species<br>(clonality) | Concentration used |  |  |
| --- | --- | --- | --- | --- | --- |
|  |  |  | Muscle | Retina | Sciatic nerve |
| Mbp | Boehringer (1118099) | Mouse<br>(M) | 1/250 | - | - |
| Nrp1 | R & D Systems (AF566) | Goat (P) | 1/100 | 1/100 | - |
| Pecam1 | Santa Cruz Biotechnology<br>(sc-1506) | Goat (P) | 1/50 | - | 1/50 |
| S100 | Dako (Z0311) | Rabbit<br>(P) | 1/200 | - | - |
| SV2 | DSHB (concentrate) | Mouse<br>(M) | 1/100 | - | - |
| Vegfr2 | CST (55B11) | Rabbit<br>(M) | 1/100 | 1/100 | - |
| 2H3 | DSHB (supernatant) | Mouse<br>(M) | 1/50 | - | - |

## References

Achilli, F., Bros-Facer, V., Williams, H. P., Banks, G. T., AlQatari, M., Chia, R., Tucci, V., Groves, M., Nickols, C. D., Seburn, K. L. et al. (2009). An ENU-induced mutation in mouse glycyl-tRNA synthetase (*GARS*) causes peripheral sensory and motor phenotypes creating a model of Charcot-Marie-Tooth type 2D peripheral neuropathy. Dis Model Mech 2, 359–73.

Antonellis, A., Ellsworth, R. E., Sambuughin, N., Puls, I., Abel, A., Lee-Lin, S. Q., Jordanova, A., Kremensky, I., Christodoulou, K., Middleton, L. T. et al. (2003). Glycyl tRNA synthetase mutations in Charcot-Marie-Tooth disease type 2D and distal spinal muscular atrophy type V. Am J Hum Genet 72, 1293–9.

Antonellis, A. and Green, E. D. (2008). The role of aminoacyl-tRNA synthetases in genetic diseases. Annu Rev Genomics Hum Genet 9, 87–107.

Bovenkamp, D. E., Goishi, K., Bahary, N., Davidson, A. J., Zhou, Y., Becker, T., Becker, C. G., Zon, L. I. and Klagsbrun, M. (2004). Expression and mapping of duplicate *neuropilin-1* and *neuropilin-2* genes in developing zebrafish. Gene Expr Patterns 4, 361–70.

Carmeliet, P., Ferreira, V., Breier, G., Pollefeyt, S., Kieckens, L., Gertsenstein, M., Fahrig, M., Vandenhoeck, A., Harpal, K., Eberhardt, C. et al. (1996). Abnormal blood vessel development and lethality in embryos lacking a single VEGF allele. Nature 380, 435–9.

Cortes, C. J., Ling, S. C., Guo, L. T., Hung, G., Tsunemi, T., Ly, L., Tokunaga, S., Lopez, E., Sopher, B. L., Bennett, C. F. et al. (2014). Muscle expression of mutant androgen receptor accounts for systemic and motor neuron disease phenotypes in spinal and bulbar muscular atrophy. Neuron 82, 295–307.

Epstein, J. A., Aghajanian, H. and Singh, M. K. (2015). Semaphorin signaling in cardiovascular development. Cell Metab 21, 163–73.

Fantin, A., Herzog, B., Mahmoud, M., Yamaji, M., Plein, A., Denti, L., Ruhrberg, C. and Zachary, I. (2014). Neuropilin 1 (NRP1) hypomorphism combined with defective VEGF-A binding reveals novel roles for NRP1 in developmental and pathological angiogenesis. Development 141, 556–62.

Fantin, A., Schwarz, Q., Davidson, K., Normando, E. M., Denti, L. and Ruhrberg, C. (2011). The cytoplasmic domain of neuropilin 1 is dispensable for angiogenesis, but promotes the spatial separation of retinal arteries and veins. Development 138, 4185–91.

Fantin, A., Vieira, J. M., Gestri, G., Denti, L., Schwarz, Q., Prykhozhij, S., Peri, F., Wilson, S. W. and Ruhrberg, C. (2010). Tissue macrophages act as cellular chaperones for vascular anastomosis downstream of VEGF-mediated endothelial tip cell induction. Blood 116, 829–40.

Fantin, A., Vieira, J. M., Plein, A., Maden, C. H. and Ruhrberg, C. (2013). The embryonic mouse hindbrain as a qualitative and quantitative model for studying the molecular and cellular mechanisms of angiogenesis. Nat Protoc 8, 418–29.

Feldner, J., Becker, T., Goishi, K., Schweitzer, J., Lee, P., Schachner, M., Klagsbrun, M. and Becker, C. G. (2005). Neuropilin-1a is involved in trunk motor axon outgrowth in embryonic zebrafish. Dev Dyn 234, 535–49.

Ferrara, N., Carver-Moore, K., Chen, H., Dowd, M., Lu, L., O’Shea, K. S., Powell-Braxton, L., Hillan, K. J. and Moore, M. W. (1996). Heterozygous embryonic lethality induced by targeted inactivation of the VEGF gene. Nature 380, 439–42.

Fujisawa, H., Takagi, S. and Hirata, T. (1995). Growth-associated expression of a membrane protein, neuropilin, in *Xenopus* optic nerve fibers. Dev Neurosci 17, 343–9.

Gelfand, M. V., Hagan, N., Tata, A., Oh, W. J., Lacoste, B., Kang, K. T., Kopycinska, J., Bischoff, J., Wang, J. H. and Gu, C. (2014). Neuropilin-1 functions as a VEGFR2 co-receptor to guide developmental angiogenesis independent of ligand binding. Elife 3, e03720.

Gerhardt, H., Golding, M., Fruttiger, M., Ruhrberg, C., Lundkvist, A., Abramsson, A., Jeltsch, M., Mitchell, C., Alitalo, K., Shima, D. et al. (2003). VEGF guides angiogenic sprouting utilizing endothelial tip cell filopodia. J Cell Biol 161, 1163–77.

Grice, S. J., Sleigh, J. N., Motley, W. W., Liu, J. L., Burgess, R. W., Talbot, K. and Cader, M. Z. (2015). Dominant, toxic gain-of-function mutations in *gars* lead to non-cell autonomous neuropathology. Hum Mol Genet 24, 4397–406.

Gu, C. and Giraudo, E. (2013). The role of semaphorins and their receptors in vascular development and cancer. Exp Cell Res 319, 1306–16.

Gu, C., Limberg, B. J., Whitaker, G. B., Perman, B., Leahy, D. J., Rosenbaum, J. S., Ginty, D. D. and Kolodkin, A. L. (2002). Characterization of neuropilin-1 structural features that confer binding to semaphorin 3A and vascular endothelial growth factor 165. J Biol Chem 277, 18069–76.

He, W., Bai, G., Zhou, H., Wei, N., White, N. M., Lauer, J., Liu, H., Shi, Y., Dumitru, C. D., Lettieri, K. et al. (2015). CMT2D neuropathy is linked to the neomorphic binding activity of glycyl-tRNA synthetase. Nature 526, 710–4.

He, W., Zhang, H. M., Chong, Y. E., Guo, M., Marshall, A. G. and Yang, X. L. (2011). Dispersed disease-causing neomorphic mutations on a single protein promote the same localized conformational opening. Proc Natl Acad Sci U S A 108, 12307–12.

Helmbrecht, M. S., Soellner, H., Truckenbrodt, A. M., Sundermeier, J., Cohrs, C., Hans, W., de Angelis, M. H., Feuchtinger, A., Aichler, M., Fouad, K. et al. (2015). Loss of Npn1 from motor neurons causes postnatal deficits independent from Sema3A signaling. Dev Biol 399, 2–14.

Huber, A. B., Kania, A., Tran, T. S., Gu, C., De Marco Garcia, N., Lieberam, I., Johnson, D., Jessell, T. M., Ginty, D. D. and Kolodkin, A. L. (2005). Distinct roles for secreted semaphorin signaling in spinal motor axon guidance. Neuron 48, 949–64.

Huettl, R. E., Soellner, H., Bianchi, E., Novitch, B. G. and Huber, A. B. (2011). Npn-1 contributes to axon-axon interactions that differentially control sensory and motor innervation of the limb. PLoS Biol 9, e1001020.

Kawasaki, T., Kitsukawa, T., Bekku, Y., Matsuda, Y., Sanbo, M., Yagi, T. and Fujisawa, H. (1999). A requirement for neuropilin-1 in embryonic vessel formation. Development 126, 4895–902.

Kofler, N. M. and Simons, M. (2015). Angiogenesis versus arteriogenesis: neuropilin 1 modulation of VEGF signaling. F1000Prime Rep 7, 26.

Lieberman, A. P., Yu, Z., Murray, S., Peralta, R., Low, A., Guo, S., Yu, X. X., Cortes, C. J., Bennett, C. F., Monia, B. P. et al. (2014). Peripheral androgen receptor gene suppression rescues disease in mouse models of spinal and bulbar muscular atrophy. Cell Rep 7, 774–84.

Lu, Q. R., Yuk, D., Alberta, J. A., Zhu, Z., Pawlitzky, I., Chan, J., McMahon, A. P., Stiles, C. D. and Rowitch, D. H. (2000). Sonic hedgehog regulated oligodendrocyte lineage genes encoding bHLH proteins in the mammalian central nervous system. Neuron 25, 317–29.

Mackenzie, F. and Ruhrberg, C. (2012). Diverse roles for VEGF-A in the nervous system. Development 139, 1371–80.

Moret, F., Renaudot, C., Bozon, M. and Castellani, V. (2007). Semaphorin and neuropilin co-expression in motoneurons sets axon sensitivity to environmental semaphorin sources during motor axon pathfinding. Development 134, 4491–501.

Motley, W. W., Seburn, K. L., Nawaz, M. H., Miers, K. E., Cheng, J., Antonellis, A., Green, E. D., Talbot, K., Yang, X. L., Fischbeck, K. H. et al. (2011). Charcot-Marie-Tooth-linked mutant GARS is toxic to peripheral neurons independent of wild-type GARS levels. PLoS Genet 7, e1002399.

Murray, L., Gillingwater, T. H. and Kothary, R. (2014). Dissection of the transversus abdominis muscle for whole-mount neuromuscular junction Analysis. J Vis Exp 83, e51162.

Nangle, L. A., Zhang, W., Xie, W., Yang, X. L. and Schimmel, P. (2007). Charcot-Marie-Tooth disease-associated mutant tRNA synthetases linked to altered dimer interface and neurite distribution defect. Proc Natl Acad Sci U S A 104, 11239–44.

Neufeld, G., Cohen, T., Shraga, N., Lange, T., Kessler, O. and Herzog, Y. (2002). The neuropilins: multifunctional semaphorin and VEGF receptors that modulate axon guidance and angiogenesis. Trends Cardiovasc Med 12, 13–9.

Oosthuyse, B., Moons, L., Storkebaum, E., Beck, H., Nuyens, D., Brusselmans, K., Van Dorpe, J., Hellings, P., Gorselink, M., Heymans, S. et al. (2001). Deletion of the hypoxia-response element in the vascular endothelial growth factor promoter causes motor neuron degeneration. Nat Genet 28, 131–8.

Pan, Q., Chanthery, Y., Liang, W. C., Stawicki, S., Mak, J., Rathore, N., Tong, R. K., Kowalski, J., Yee, S. F., Pacheco, G. et al. (2007). Blocking neuropilin-1 function has an additive effect with anti-VEGF to inhibit tumor growth. Cancer Cell 11, 53–67.

Park, M. C., Kang, T., Jin, D., Han, J. M., Kim, S. B., Park, Y. J., Cho, K., Park, Y. W., Guo, M., He, W. et al. (2012). Secreted human glycyl-tRNA synthetase implicated in defense against ERK-activated tumorigenesis. Proc Natl Acad Sci U S A 109, E640–7.

Pitulescu, M. E., Schmidt, I., Benedito, R. and Adams, R. H. (2010). Inducible gene targeting in the neonatal vasculature and analysis of retinal angiogenesis in mice. Nat Protoc 5, 1518–34.

Plein, A., Fantin, A. and Ruhrberg, C. (2014). Neuropilin regulation of angiogenesis, arteriogenesis, and vascular permeability. Microcirculation 21, 315–23.

Puentes, F., Malaspina, A., van Noort, J. M. and Amor, S. (2016). Non-neuronal Cells in ALS: Role of Glial, Immune cells and Blood-CNS Barriers. Brain Pathol 26, 248–57.

Ruhrberg, C., Gerhardt, H., Golding, M., Watson, R., Ioannidou, S., Fujisawa, H., Betsholtz, C. and Shima, D. T. (2002). Spatially restricted patterning cues provided by heparin-binding VEGF-A control blood vessel branching morphogenesis. Genes Dev 16, 2684–98.

Ruiz de Almodovar, C., Lambrechts, D., Mazzone, M. and Carmeliet, P. (2009). Role and therapeutic potential of VEGF in the nervous system. Physiol Rev 89, 607–48.

Schwarz, Q., Gu, C., Fujisawa, H., Sabelko, K., Gertsenstein, M., Nagy, A., Taniguchi, M., Kolodkin, A. L., Ginty, D. D., Shima, D. T. et al. (2004). Vascular endothelial growth factor controls neuronal migration and cooperates with Sema3A to pattern distinct compartments of the facial nerve. Genes Dev 18, 2822–34.

Seburn, K. L., Nangle, L. A., Cox, G. A., Schimmel, P. and Burgess, R. W. (2006). An active dominant mutation of glycyl-tRNA synthetase causes neuropathy in a Charcot-Marie-Tooth 2D mouse model. Neuron 51, 715–26.

Shalaby, F., Rossant, J., Yamaguchi, T. P., Gertsenstein, M., Wu, X. F., Breitman, M. L. and Schuh, A. C. (1995). Failure of blood-island formation and vasculogenesis in Flk-1-deficient mice. Nature 376, 62–6.

Sleigh, J. N., Burgess, R. W., Gillingwater, T. H. and Cader, M. Z. (2014a). Morphological analysis of neuromuscular junction development and degeneration in rodent lumbrical muscles. J Neurosci Methods 227, 159–65.

Sleigh, J. N., Dawes, J. M., West, S. J., Spaulding, E. L., Gomez-Martin, A., Burgess, R. W., Cader, M. Z., Talbot, K., Bennett, D. L. and Schiavo, G. (2016). Sensory neuron fate is developmentally perturbed by *Gars* mutations causing human neuropathy. BioRxiv doi:http://dx.doi.org/10.1101/071159.

Sleigh, J. N., Gillingwater, T. H. and Talbot, K. (2011). The contribution of mouse models to understanding the pathogenesis of spinal muscular atrophy. Dis Model Mech 4, 457–67.

Sleigh, J. N., Grice, S. J., Burgess, R. W., Talbot, K. and Cader, M. Z. (2014b). Neuromuscular junction maturation defects precede impaired lower motor neuron connectivity in Charcot-Marie-Tooth type 2D mice. Hum Mol Genet 23, 2639–50.

Soker, S., Takashima, S., Miao, H. Q., Neufeld, G. and Klagsbrun, M. (1998). Neuropilin-1 is expressed by endothelial and tumor cells as an isoform-specific receptor for vascular endothelial growth factor. Cell 92, 735–45.

Somers, E., Lees, R. D., Hoban, K., Sleigh, J. N., Zhou, H., Muntoni, F., Talbot, K., Gillingwater, T. H. and Parson, S. H. (2016). Vascular defects and spinal cord hypoxia in spinal muscular atrophy. Ann Neurol 79, 217–30.

Somers, E., Stencel, Z., Wishart, T. M., Gillingwater, T. H. and Parson, S. H. (2012). Density, calibre and ramification of muscle capillaries are altered in a mouse model of severe spinal muscular atrophy. Neuromuscul Disord 22, 435–42.

Spaulding, E. L., Sleigh, J. N., Morelli, K. H., Pinter, M. J., Burgess, R. W. and Seburn, K. L. (2016). Synaptic deficits at neuromuscular junctions in two mouse models of Charcot-Marie-Tooth type 2d. J Neurosci 36, 3254–67.

Tata, M., Ruhrberg, C. and Fantin, A. (2015). Vascularisation of the central nervous system. Mech Dev 138 Pt 1, 26–36.

Venkova, K., Christov, A., Kamaluddin, Z., Kobalka, P., Siddiqui, S. and Hensley, K. (2014). Semaphorin 3A signaling through neuropilin-1 is an early trigger for distal axonopathy in the SOD1G93A mouse model of amyotrophic lateral sclerosis. J Neuropathol Exp Neurol 73, 702–13.

Xie, W., Nangle, L. A., Zhang, W., Schimmel, P. and Yang, X. L. (2007). Long-range structural effects of a Charcot-Marie-Tooth disease-causing mutation in human glycyl-tRNA synthetase. Proc Natl Acad Sci U S A 104, 9976–81.

